# Speech tracking in auditory and motor regions reflects distinct linguistic features

**DOI:** 10.1101/195941

**Authors:** Anne Keitel, Joachim Gross, Christoph Kayser

## Abstract

During online speech processing, our brain tracks the acoustic fluctuations in speech at different time-scales. Previous research has focussed on generic time-scales (for example, delta or theta bands) that are assumed to map onto linguistic features such as prosody or syllables. However, given the high inter-subject variability in speaking patterns, such a generic association between the time-scales of brain activity and speech properties can be ambiguous. Here, we analyse speech tracking in source-localised magnetoencephalographic data by directly focusing on time-scales extracted from statistical regularities in the speech material. This revealed widespread tracking at the time-scales of phrases (0.6 – 1.3 Hz), words (1.8 – 3 Hz), syllables (2.8 – 4.8 Hz), and phonemes (8 – 12.4 Hz). Importantly, when examining the relevance for single-trial comprehension, we found stronger tracking for correctly comprehended trials in the left premotor cortex at the phrasal scale, and in left middle temporal cortex at the word scale. Control analyses using generic bands confirmed that these effects were specific to the stimulus-tailored speech regularities. Furthermore, we found that the phase at the phrasal time-scale coupled to beta-power in motor areas. This cross-frequency coupling likely mediates the comprehension effect in the motor system, and implies top-down temporal prediction in speech perception. Together, our results reveal specific functional and perceptually relevant roles of distinct entrainment processes along the auditory-motor pathway. These processes act concurrently at time-scales within the traditional delta band and highlight the role of neural tracking mechanisms that reflect the temporal characteristics of speech.

---

Speech consists of hierarchically organised linguistic segments such as phrases, words, syllables, and phonemes. To comprehend speech, a listener needs to parse the continuous stream into these segments. One mechanism that has been proposed to fulfil such a role is the tracking of speech information in dynamic brain activity (often termed speech-to-brain entrainment) [1,2]. Such tracking is observed as a precise alignment of rhythmic brain activity to the temporal characteristics of speech. Previous studies commonly focussed on brain activity at the time-scales of traditional delta (1 – 4 Hz) and theta (4 – 8 Hz) bands [3,4,5,6]. These have been suggested to reflect the neural processing of prosodic and syllabic speech features. Accordingly, when looking at the speech signal itself these features often emerge at time-scales matching the generic delta and theta bands [7,8]. However, such a general association between time-scales in brain activity and speech properties is difficult [9]. First, there are large inter-individual differences in speaking rate and use of prosody. Second, the association between specific time-scales and linguistic or phonological features remains vague as multiple properties have been associated with delta (stress, intonation, phrase-structure) [10,11,12] and theta bands (syllables, jaw movements) [13,14]. In the present study, we thus took a different approach and first extracted linguistic time-scales from the speech corpus, based on stimulus-specific regularities. Effects at these linguistic time-scales are easier to interpret than effects at generic time-scales, and this approach thereby helps to gain specificity about tracking processes.

The functional interpretation of speech-to-brain entrainment is further hampered by a lack of a systematic assessment of where (in the brain) and at which time-scale it is relevant for comprehension. Most studies attempting to link comprehension and speech tracking have done so only indirectly [but see, 15], by modulating speech intelligibility using background noise [16,17], noise vocoding [5,18], or reversed speech [3,19,20]. These studies revealed a link between delta and theta band tracking and intelligibility, by relating neural markers to acoustic speech properties during listening to continuous speech [2,3,15]. Some of these studies have also demonstrated widespread low-frequency tracking across the brain, going beyond auditory cortex and encompassing prefrontal and motor areas [3,16]. However, it remains uncertain whether and where the tracking of speech by dynamic brain activity is necessary for comprehension at the single trial level.

In the present study, participants performed a comprehension task on natural, structurally identical sentences embedded in noise. Sentences consisted of a set of words that were randomly combined and therefore semantically unpredictable, but meaningful. Using magnetoencephalography (MEG) we analysed speech tracking (quantified by mutual information, MI) in source-localised data at the single trial level. We then directly tested the perceptual relevance of speech tracking at time-scales mapping onto linguistic categories such as phrasal elements, words, syllables and phonemes.

## Results

### Behavioural results

Participants listened to single sentences and indicated after each trial which (out of four) target words occurred in the sentence. Participants reported the correct target word on average in 69.7 ± 7.1% (*M* ± *SD*) of trials, with chance level being 25%. Performance of individual participants ranged from 56.1% to 81.1%, allowing a comparison between correct and incorrect trials for each participant.

### Speech tracking across all trials

The mutual information (MI) between the source-localised, Hilbert-transformed MEG time-series and the Hilbert-transformed speech envelope was computed within four frequency bands. These reflected the rates of phrases (0.6 – 1.3 Hz), words (1.8 – 3 Hz), syllables (2.8 – 4.8 Hz), and phonemes (8 – 12.4 Hz) in the stimulus corpus. The boundaries of each band were defined based on the slowest and fastest event per linguistic category across sentences (see Methods). When compared to surrogate data, speech MI was significant in all analysed frequency bands (see supplemental **Figure S1**). As in previous studies, MI was strongest in early auditory areas [3,16], and decreased with increasing frequency [4,16]. Tracking of phrases, words, and syllables was reflected in a bilateral cluster, whereas phoneme-tracking was only significant in the right hemisphere

### Perceptually relevant speech tracking

To localise cortical regions where entrainment was relevant for comprehension, we statistically compared MI between correct and incorrect trials within each band. This yielded significant left-hemispheric clusters in two frequency bands (**Figure 1B**). For the phrasal time-scale (0.6 – 1.3 Hz), MI was larger for correctly vs. incorrectly comprehended trials in a cluster comprising left pre- and postcentral regions, supramarginal gyrus, and Heschl gyrus (*T*_sum_(19) = 568.00, *p*_cluster_ = .045, 205 grid points). The effect peaked in the left premotor cortex (PM, left BA 6). For the word time-scale (1.8 – 3 Hz), MI was larger in a cluster comprising left superior, middle, and inferior temporal gyrus as well as supramarginal gyrus and Heschl gyrus (*T*_sum_(19) = 739.59, *p*_cluster_ = .018, 263 grid points). The effect peaked in the left middle temporal gyrus (MTG, left BA 21). There was a small overlap of the clusters for phrasal and word time-scales, peaking in the left Heschl gyrus (HG, left BA 41).

**Figure 1.**
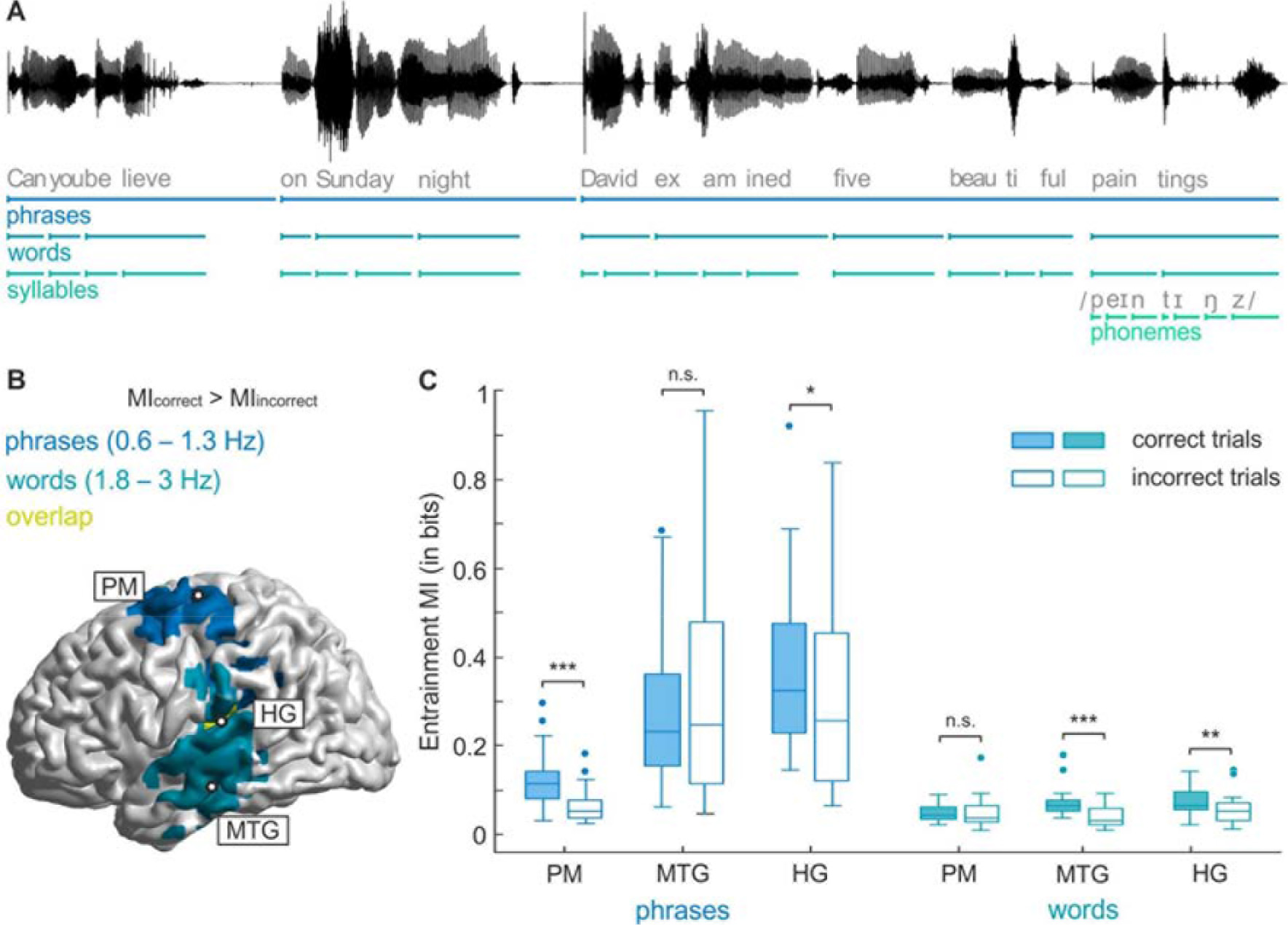
Stimulus material and main results. **A)** Example sentence from the stimulus material. Shown is the acoustic waveform (black line) as well as its segmentation into phrases, words, syllables, and phonemes (last word only). **B)** Brain regions relevant for comprehension. Clusters where speech MI was larger for correctly compared to incorrectly comprehended trials (*t*-test, cluster-corrected). For phrases (blue), the effect was strongest in the left premotor (PM) area. For words (turquoise), it was strongest in the left middle temporal gyrus (MTG). Areas that were perceptually relevant for both phrases and words (yellow) include left Heschl gyrus (HG) and supramarginal gyrus. Peak grid points are denoted with circles. **C)** Comparison of MI values at peak grid points in premotor gyrus (PM), middle temporal gyrus (MTG), and Heschl gyrus (HG). Boxes denote interquartile range with median line, error bars show minimum and maximum, excluding outliers. Significance is denoted with: ***: *p* < .001, **: *p* < .01, *: *p* < .05, n.s.: not significant.

To determine whether these effects are specific to those time-scales extracted from the stimulus corpus, we performed several control analyses. First, we performed post-hoc *t*-tests at the peak grid points of each cluster to see whether phrasal and word effects are indeed significant only for the respective time-scales (**Figure 1C**). As expected, MI differed between correct and incorrect trials at the phrasal time-scale in left premotor cortex and Heschl gyrus (*t*(19) = 4.90, *p*_FDR_ < .001 and *t*(19) = 2.53, *p*_FDR_ = .031, respectively). Likewise, MI differed at the word time-scale in the left middle temporal gyrus and Heschl gyrus (*t*(19) = 5.22, *p*_FDR_ < .001 and *t*(19) = 3.48, *p*_FDR_ = .005, respectively). Importantly, MI neither differed at the phrasal scale in MTG (*t*(19) = −1.78, *p*_FDR_ = .11), nor at the word scale in premotor gyrus (*t*(19) = −0.57, *p*_FDR_ = .58).

Second, we also compared MI between correctly and incorrectly comprehended trials in seven generic, 2-Hz wide frequency bands (from 0 – 8 Hz, in 1-Hz steps) to confirm that the above intelligibility-related effects are indeed specific to those frequency bands matched to the specific temporal structure of the speech material. This is an important control, as most previous studies use generic bands with a predefined fixed frequency spacing. Significant comprehension effects were found in two bands (supplemental **Figure S2A**). For the 1 – 3 Hz band, MI was larger for comprehended than uncomprehended trials in a cluster centred around auditory cortex (*T*_sum_(19) = 1,078.85, *p*_ciuster_ = .030). For the 2 - 4 Hz band, MI was larger in a cluster covering middle and inferior temporal cortex (*T*_sum_(19) = 751.93, p_cluster_ = .046).

Third, using post-hoc statistics, we also verified that MI at the previously identified peak grid points (see **Figure 1B**) differed between correct and incorrect trials only in those sub-bands that match the stimulus-specific time-scales (supplementary **Figure S2B**). Notably, the motor cortex was not found to mediate a comprehension effect in any of the generic bands.

### Phase-amplitude coupling in the motor cortex

Rhythmic brain activity represents neuronal excitability changes [21]. In auditory areas, this mechanism has been suggested to reflect a segmentation of the incoming sensory stream [22,23]. But what is the role of slow excitability changes in the motor system? The motor system plays a role in the temporal prediction of rhythms and beats [24,25,26,27]. Previous studies have suggested that these predictions rely on the coupling of delta-phase to rhythmic activity in the beta band [24,28,29]. We therefore hypothesised that speech entrainment at the phrasal scale, and its perceptual relevance, is directly linked to phase-amplitude coupling with motor cortical beta activity. Indeed, we found that the coupling of the phase of phrasal-scale activity (0.6 – 1.3 Hz) to beta power was significantly stronger for correctly vs. incorrectly comprehended trials in our motor cluster (*t*(19) = 2.96, *p*_FDR_ = .032; see **Figure 2A**). In contrast, there was no such cross-frequency coupling for the phase of word-scale activity relative to beta power (*t*(19) = 1.14, *p*_FDR_ = .356), or of phrasal-phase to either alpha (*t*(19) = 1.38, *p*_FDR_ = .356) or theta power (*t*(19) = −0.38, *p*_FDR_ = .708).

**Figure 2.**
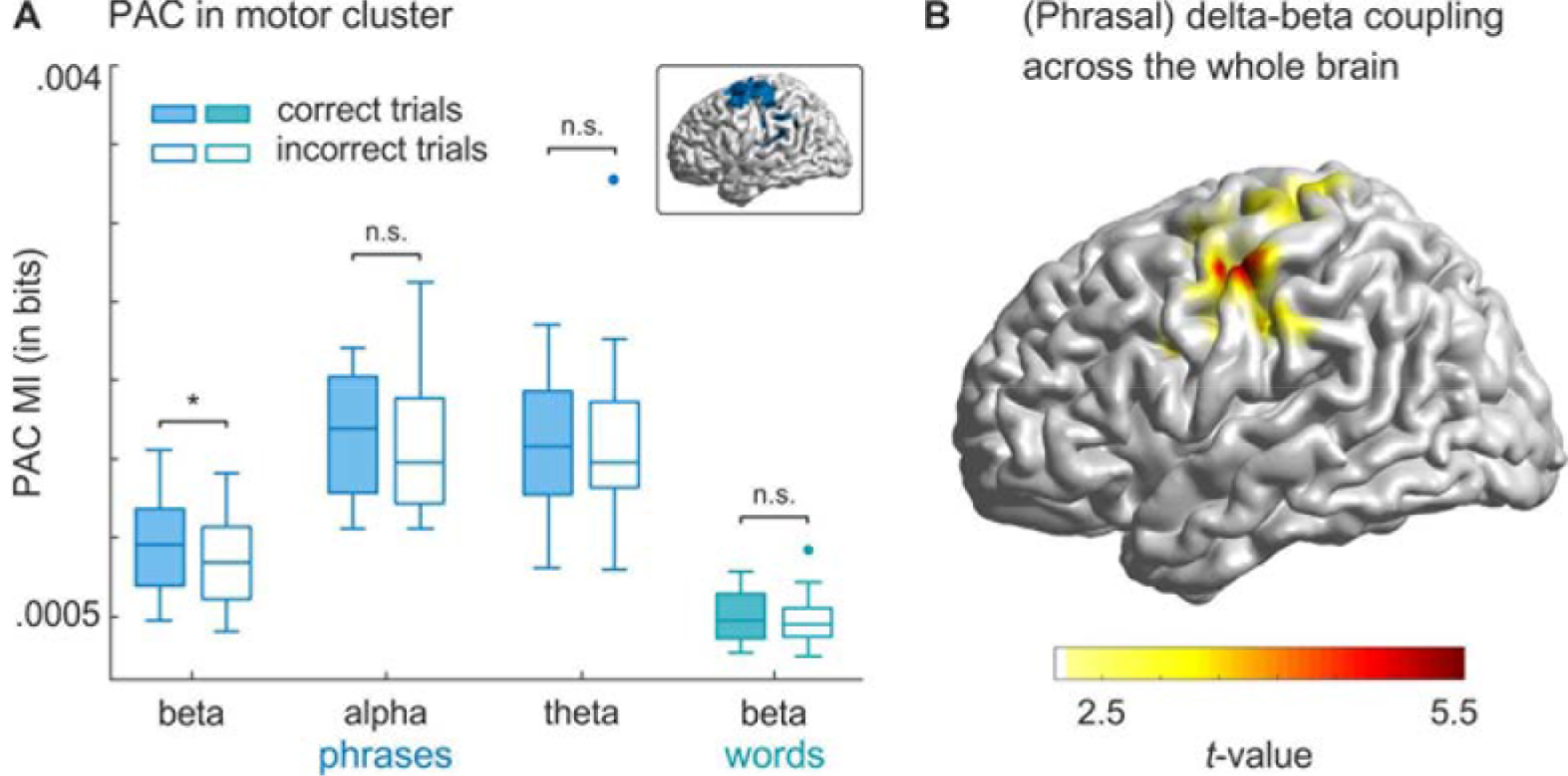
Phase-amplitude coupling (PAC) in the motor cluster and across the whole brain. A) MI between the phase at the phrasal time-scale (0.6 – 1.3 Hz) and power in beta, alpha, and theta bands (blue plots), as well as the phase at the word time-scale (1.8 – 3 Hz) and beta-power (turquoise plot) for correctly and incorrectly comprehended trials, averaged across all grid points in the motor cluster (cluster shown in inlet). Only PAC between the phase at the phrasal time-scale and beta power showed a comprehension effect. Significance is denoted with: *: *p* < .05, n.s.: not significant. B) Whole-brain analysis across all 12,337 grid points confirming that PAC between phrasal phase and beta-power is indeed confined to motor regions. Coloured area denotes the cluster where PAC between phrasal phase and beta power was larger for correctly comprehended than uncomprehended trials.

As a further control analysis, and to explore the spatial extent of PAC between phrasal phase and beta-power, we performed a whole brain analysis. We again compared PAC between correct and incorrect trials. This analysis yielded one cluster where PAC was larger for comprehended than uncomprehended trials (*T*_sum_ = 203.94, *p*_cluster_ = .030, one-sided; see **Figure 2B**). The cluster comprised left pre- and postcentral regions. Thus, behaviourally relevant PAC was confined to the left motor system, overlapping with the speech tracking effect in left motor areas.

## Discussion

By focusing on stimulus-specific time-scales and measuring comprehension on individual trials, we show that distinct neural and linguistic time-scales relate to speech encoding along the auditory-motor pathway. Our findings provide specific functional roles of speech entrainment at two time-scales within the delta band, one relevant in motor areas, accompanied by modulations in delta-beta coupled oscillations, and one relevant in temporal areas.

### The motor system predicts phrasal timing using beta oscillations

We show that speech tracking at the phrasal time-scale is relevant for comprehension in left motor areas. The premotor cortex, and the motor system generally, has been associated with generating temporal predictions [30,31,32,33,34] and the processing of rhythms and beats [25,26,27]. In the current stimulus corpus, the timing of phrasal elements was relatively predictable, as sentences followed the same structure. The phrasal structure was also defined by prominent pauses between phrasal elements (evident in the clear peak in the frequency spectrum, supplementary **Figure S3A**). On the other hand, words (and therefore syllables and phonemes) were not semantically, or temporally, predictable due to the scrambling of words across sentences. The motor system likely exploited the temporally predictive phrasal information for parsing and segmenting the sentences, therefore facilitating comprehension. Yet, it is possible that this mechanism is not specific to the phrasal structure per se. Instead, it could be that the motor system would exploit *any* temporal regularities [26], regardless of their linguistic or meta-linguistic relevance. Future research is required to directly compare acoustic and linguistic regularities and their relevance for speech tracking.

It has been hypothesised that delta-entrainment to speech in the left hemisphere reflects a motor-driven top-down modulation [35]. These top-down modulations have been associated with beta oscillations [31,36,37,38], which are prevalent in the motor system [39]. The finding that the temporal prediction of tone sequences is mediated by pre-stimulus delta-beta coupled oscillations further supports this hypothesis [24,see also 28,29]. Here, we show, for the first time, to our knowledge, that such a cross-frequency mechanism can also operate during speech perception. This coupling is a) specific to phrasal delta-phase (0.6 – 1.3 Hz) and beta-power (13 – 30 Hz), and b) only behaviourally relevant in left motor areas. Our results thereby underpin a mechanistic role of delta-beta coupling in the motor system for speech comprehension.

### Word segmentation in the temporal cortex

Speech tracking at the word-scale was perceptually relevant across the entire mid-temporal gyrus, peaking in middle temporal gyrus (MTG) and including superior and inferior temporal gyrus, and inferior supramarginal gyrus. The MTG is associated with lexical semantic processes [40,41], and is one end point of the ventral auditory pathway, mapping sound-to-meaning [42]. It is plausible that stronger speech tracking, and therefore better word-scale segmentation in these regions is directly linked to comprehension performance. The result that the comprehension effect at the word-scale extends dorsally to supramarginal gyrus seems to contradict models of a ventral focus of word comprehension. However, it is consistent with the notion of a dorsal lexicon, thought to store articulatorily organised word form representations [43].

### Specificity of linguistic time-scales

An analysis of 2-Hz wide generic bands showed that 1) the motor system was not predictive for comprehension in any generic band, 2) the 1 – 3 Hz band yielded a similar pattern as the word time-scale with which it overlaps, and 3) the 2 – 4 Hz band also overlapped with the effect at the word time-scale, albeit less so (supplemental **Figure S2**). These results suggest that perceptually relevant speech tracking in the motor system is specific to the phrasal time-scale in the stimulus material. On the other hand, a comprehension effect in temporal regions is found in the delta band (above 1 Hz and below 4 Hz), independent of the specific boundaries of the used bands (although 0 – 2 Hz did not yield a significant effect). This suggests that speech tracking in temporal areas emerges at more widespread time-scales, perhaps because word length is more variable than phrasal length in the present stimulus material. Analyses of the coefficient of variation supported this interpretation: When compared with phrases (*c*_v_ = 0.27), words varied in length almost twice as much (*c*_v_ = 0.48).

We chose to base the time-scales on linguistic categories of phrases, words, syllables and phonemes. This is the most pragmatic approach, as the language system ultimately has to parse the speech stream into these segments. However, one could argue that these linguistic categories overlap with other meta-linguistic elements that also follow temporal modulations below 4 Hz such as prosodic features [12]. The most relevant prosodic features for speech segmentation are pauses, stress, and intonation [10,44]. The phrases in the stimulus material are *defined by* pauses and therefore phrasal time-scale and timing of pauses can be considered one and the same. The interaction between linguistic categories and lexical stress is more complex. If we consider every third syllable stressed [45], we yield a “stress time-scale” of 0.9 – 1.6 Hz, which partly overlaps with the phrasal time-scale (0.6 – 1.3 Hz). The role of stress is manifold in speech (disambiguation of phonemically identical words, highlighting the meaning of words, metrical stress) and common sense would suggest that it makes little sense to segment the speech stream by using stressed syllables as boundaries. Thus, although stress is important and useful in speech comprehension, focussing on the phrasal time-scale (as opposed to the “stress time-scale”) is more appropriate. Fluctuations in pitch, or intonation, also occur in the delta band (see supplementary **Figure S3B** for spectral analysis of pitch, or its acoustic correlate the fundamental frequency). Pitch fluctuations can signal phrasal boundaries [46], and an overlap with the phrasal time-scale is therefore not surprising. As the auditory system is able to track pitch fluctuations [9], and fundamental frequency and intensity are related, we cannot completely disentangle pitch tracking from envelope tracking. But language comprehension requires the grouping of words into phrases [11], and focussing the analysis on the phrasal time-scale is the most direct way of analysing phrasal processing. Future research needs to address the question to what extent phrasal segmentation relies on the acoustic envelope, pitch fluctuations, or both.

Taken together, linguistic and meta-linguistic events in natural speech have a tendency to co-occur [47] and their interaction is complex. However, for natural speech processing, the division into linguistic categories, as done in the present study, seems the most pragmatic and ecologically valid solution to gain specificity about speech comprehension effects.

In the present study, the average speech rate was ~130 words/minute. In two other studies that reported speech rate, it was considerably higher, at ~160 words/minute [16] and ~210 words/minute [19]. The rate of syllables is typically associated with frequencies between 4 and 8 Hz [7,48]. In the present study it was 2.8 – 4.8 Hz, and in another study it was even lower at 2 – 4 Hz [1]. These differences demonstrate that, even in experimental contexts, speech rates can deviate from the assumed standard. Furthermore, a recent study has shown that the auditory system is not limited by traditionally imposed frequency bands [36]. It therefore is highly beneficial to calculate stimulus-specific speech statistics for speech tracking analyses, instead of applying generic frequency bands.

### Not all speech tracking processes are relevant for comprehension

Our results regarding overall speech tracking (compared with surrogate data) replicate previous reports of wide-spread speech-to-brain entrainment at multiple time-scales [3,16]. However, in our data only speech tracking in specific bands within the delta frequency range was directly relevant for perception. This is in line with different functional roles for speech tracking below and above 3 Hz [49]. In particular, speech tracking at the syllabic and phonetic scales did not differ between trials with correct and incorrect comprehension. Although, for the participants to comprehend target words correctly, at least some syllables must have been encoded phonetically. It could be that the use of a noisy background prevented the robust encoding of individual syllables or phonemes, therefore reducing the tracking at these time-scales or reducing the statistical power in detecting between-trial differences. However, auditory entrainment also occurs for sub-threshold stimuli [50], suggesting that under some circumstances the presence of entrainment per se is not a sufficient marker for the relevant processes to influence perception. Additional work is required to understand whether speech tracking at the syllabic and phonetic time-scales is indeed a robust marker of the actual neural encoding of these features, or whether only speech tracking at time-scales below the syllabic rate directly indexes functionally and perceptually relevant processes. Furthermore, the left-lateralised comprehension effects at slow time-scales stand in contrast with bilateral overall speech tracking at these scales (supplemental **Figure S1**). This supports the notion that “early” acoustic processes are bilateral, whereas “higher-order” speech comprehension is left-lateralised [51].

## Materials and Methods

### Participants and data acquisition

Following previous sample sizes of MEG-studies that used mutual information to study speech tracking [3,16], 20 healthy, native British participants took part in the study (9 female, age 23.6 ± 5.8 years [*M* ± *SD*]). All participants were right-handed [Edinburgh Handedness Inventory; 52], had normal hearing [Quick Hearing Check; 53] and normal or corrected-to-normal vision. Furthermore, participants had no self-reported current or previous neurological or language disorders. All participants provided written informed consent prior to testing and received monetary compensation of £10/h. The experiment was approved by a local ethics committee (College of Science and Engineering, University of Glasgow), and conducted in compliance with the Declaration of Helsinki.

MEG was recorded with a 248-magnetometers, whole-head MEG system (MAGNES 3600 WH, 4-D Neuroimaging) at a sampling rate of 1 KHz. Head positions were measured at the beginning and end of each run, using five coils placed on the participants’ head. Coil positions were co-digitised with head-shape (FASTRAK®, Polhemus Inc., VT, USA). Participants sat upright and fixated a fixation point projected centrally on screen with a DLP projector. Sounds were transmitted binaurally through plastic earpieces and 5-m long plastic tubes connected to a sound pressure transducer. Stimulus presentation was controlled with Psychophysics toolbox [54] for MATLAB (The MathWorks, Inc.).

### Stimuli

The stimulus material consisted of two equivalent sets of 90 sentences (180 in total) that were spoken by a trained, male, native British actor. The speaker was instructed to speak clearly and naturally. Sentences were constructed to be meaningful, but unpredictable. Each sentence consisted of the same basic elements and therefore had the same structure. For example, the sentence “[Did you notice], [on Sunday night], [Graham] [offered] [ten] [fantastic] [books]”, consists of a “filler” phrase, a time phrase, a name, a verb, a number, an adjective, and a noun. There were 18 possible names, verbs, numbers, adjectives, and nouns that were each repeated ten times. Sentence elements were randomly combined within a set of 90 sentences. To measure comprehension, a target word was included that was either the adjective in one set of sentences (‘fantastic’ in the above example, or ‘beautiful’ in **Figure 1A**) or the number in the other set (for example, ‘twenty-one’). The duration of sentences ranged from 4.2 s to 6.5 s (5.4 ± 0.4 s [*M* ± *SD*]). Sentences were presented at a sampling rate of 22,050 Hz.

During the experiment, speech stimuli were embedded in noise. The noise consisted of ecologically valid, environmental sounds (traffic, car horns, people talking), combined into a mixture of 50 different background noises. The individual noise level for each participant was determined with a staircase procedure that was designed to yield a performance of around 70% correct. For the staircase procedure, only the 18 possible target words were used instead of whole sentences. Participants were presented with single target words embedded in noise and subsequently saw two alternatives on screen. They indicated by button press which word they had heard. Depending on whether their choice was correct or incorrect, the noise level was increased or decreased (one-up-three-down procedure) until a reliable level was reached. The average signal-to-noise ratio across participants was approximately −6 dB.

### Experimental design

The 180 sentences were presented in four blocks with 45 sentences each. In each block, participants either indicated the comprehended adjective or the comprehended number, resulting in two ‘adjective blocks’ and two ‘number blocks’. The order of sentences and blocks was randomised for each participant. The first trial of each block was a ‘dummy’ trial that was discarded for subsequent analysis; this trial was repeated at the end of the block.

After each sentence, participants were prompted with four target words (either adjectives or numbers) on the screen. They then had to indicate which one they heard by pressing one of four buttons on a button box. After 2 seconds, the next trial started automatically.

Each block lasted approximately 10 minutes, and participants could rest in between blocks. The session, including instructions, questionnaires, preparation, staircase procedure, and four blocks, took approximately 3 to 3.5 hours.

### Speech pre-processing

For each sentence, we computed the wideband speech envelope at a sampling rate of 150 Hz following procedures of previous studies [3,4,55,56]. Acoustic waveforms were first filtered into eight frequency bands (between 100 – 8,000 Hz; third order Butterworth filter; forward and reverse) that were equidistant on the cochlear frequency map [56]. From these eight individual bands, the wideband speech envelope was extracted by averaging the magnitude of the Hilbert transformed signals from each band.

To define the time-scales on which to probe speech encoding, we evaluated the rates of phrases, words, syllables, and phonemes in the stimulus material. For this, the duration between onsets of linguistic categories (here: phrases, words, and phonemes) was calculated. The exact onset timing was extracted from the speech signals using Penn Phonetics Lab Forced Aligner [P2FA; 57]. Phrases were defined as the first two clauses in each sentence (for example, [I have heard] and [on Tuesday night]). These phrases had distinct pauses (see **Fig. 1A** for an example sentence) that determined the rhythm of the sentence (also visible in the frequency spectrum in supplemental **Figure S3A**). The syllable rate is generally difficult to assess [14,58]. Here, we chose to count the actually produced syllables for each sentence. Finally, time-scales were converted to frequencies and the specific frequency bands for each category were then defined as the minimum and maximum frequencies across all 180 sentences. This led to the following bands: 0.6 – 1.3 Hz (phrases), 1.8 – 3.0 Hz (words), 2.8 – 4.8 Hz (syllables), and 8 – 12.4 Hz (phonemes). Mean and standard deviations for linguistic categories were as follows: 1.0 ± 0.1 Hz for phrases, 2.4 ± 0.3 Hz for words, 3.8 ± 0.4 Hz for syllables, and 10.4 ± 0.8 Hz for phonemes.

### MEG pre-processing

Pre-processing of MEG data was carried out in MATLAB (The MathWorks, Inc) using the Fieldtrip toolbox [59]. The four experimental blocks were pre-processed separately. Single trials were extracted from continuous data starting 2 sec before sound onset and until 10 sec after sound onset. MEG data were denoised using the reference signal. Known faulty channels (*N* = 7) were removed before further pre-processing. Trials with SQUID jumps (3.5% of trials) were detected and removed using Fieldtrip procedures with a cutoff *z*-value of 30. Before further artifact rejection, data were filtered between 0.2 and 150 Hz (fourth order Butterworth filters, forward and reverse) and down-sampled to 300 Hz. Data were visually inspected to find noisy channels (4.37 ± 3.38 on average across blocks and participants) and trials (0.66 ± 1.03 on average across blocks and participants). Finally, heart and eye movement artifacts were removed by performing an independent component analysis with 30 principal components. Data were further down-sampled to 150 Hz to match the sampling rate of the speech signal.

### Source localisation

Source localisation was performed using Fieldtrip, SPM8, and the Freesurfer toolbox. We acquired T1-weighted structural magnetic resonance images (MRIs) for each participant. These were co-registered to the MEG coordinate system using a semi-automatic procedure [3,4]. MRIs were then segmented and linearly normalised to a template brain (MNI space). A volume conduction model was constructed using a single-shell model [60]. We projected sensor-level waveforms into source space using frequency specific linear constraint minimum variance (LCMV) beamformers [61] with a regularisation parameter of 7% and optimal dipole orientation (singular value decomposition method). Grid points had a spacing of 6 mm, resulting in 12,337 points covering the whole brain.

### Analysis of speech tracking in brain activity

We quantified the statistical dependency between the speech envelope and the source-localised MEG data using mutual information (MI) [3,4,22,62]. The speech envelopes, as well as MEG data, were filtered in the four frequency bands reflecting the rates of each linguistic category using third order (for delta and theta bands) forward and reverse Butterworth filters. Within these bands, we computed the Hilbert transform and used real and imaginary parts for further analysis. Both parts were normalised separately and combined as a two-dimensional variable for the MI calculation [62]. To take into account the stimulus-brain lag, we computed MI at five different lags (from 60 to 140 ms in 20-ms steps) and summed the MI values across lags. This procedure prevents spurious results that can occur when using a single lag. First, we calculated the overall MI for each source grid point. For a robust computation of MI values, we concatenated MEG and speech data from all trials. The resulting MI values were compared with surrogate data to determine their statistical significance. Surrogate data were created by randomly shuffling trials 50 times and averaging surrogate MI values across iterations. We used a dependent *t*-test for statistical comparison for each grid point and corrected for multiple comparisons with cluster-based permutation. Specifically, we used Monte-Carlo randomisation with 1000 permutations and a critical *t*-value of 2.1, which represents the critical value of the Student’s *t*-distribution for 20 participants and a two-tailed probability of *p* = .05. The significance level for accepting clusters was 5%.

For the analysis of behavioural relevance, we compared MI between trials on which participants responded correctly and incorrectly. As the number of trials differed between these groups, we performed the calculations based on 80% of trials of the smaller sample. Again, trials were concatenated to yield robust MI values. We calculated MI values for 20 random samples of correct / incorrect trials and averaged the resulting values. MI values between correct and incorrect trials were compared using the same method and parameters as for the comparison between overall MI and surrogate MI.

To examine the specificity of the effects, we compared MI between correct and incorrect trials for all peak grid points in both frequency bands (i.e., phrasal and word time-scales). Peak grid points were those with the largest *t*-values in each cluster, and the largest summed *t*-values for the overlap of grid points. This led to six comparisons (three different grid points × two frequency bands). MI values were compared using dependent samples *t*-tests, corrected for multiple comparisons using the FDR method [63].

### Phase-amplitude coupling

To examine the hypothesis that beta-power is coupled with delta-phase in the motor cluster and that this is related to comprehension, we quantified phase-amplitude coupling (PAC) using the mutual information between beta-power and delta-phase. Phase and power were derived from Hilbert-transformed time series, and filtered in the phrasal (0.6 – 1.3 Hz) and beta band (13 – 30 Hz). Phase was expressed as a unit magnitude complex number. To get an equal number of trials for correct and incorrect trials, we again took 80% of trials of the smaller sample, concatenated trials, and repeated the calculation 50 times. This was done for all grid points within the motor cluster (*N* = 205), and then averaged across grid points and iterations. PAC was compared between correct and incorrect trials across participants using a dependent sample *t*-test.

We performed three control analyses within the motor cluster to address the frequency specificity of the effect. First, we analysed PAC between phrasal phase (0.6 – 1.3 Hz) and alpha-power (8 – 12 Hz) as well as theta-power (4 – 8 Hz). Second, we analysed PAC between the *word* phase (1.8 – 3 Hz) and beta-power. All *p*-values were corrected for multiple comparisons using the FDR method [63].

To address the spatial specificity of the delta-beta PAC coupling, we also performed a whole-brain analysis. Based on the results in the motor cluster, we hypothesised that PAC should be larger in correct than incorrect trials. PAC between phrasal delta-phase (0.6 – 1.3 Hz) and beta-power (13 – 30 Hz) was compared between correct and incorrect trials, again equalling sample sizes by using 80% of trials of the smaller sample and repeating the analysis 20 times. PAC MI was averaged across all iterations and then compared between correct and incorrect trials across participants using a dependent sample *t*-test for each grid point. To correct for multiple comparisons, we used the same parameters for cluster correction as in all previous analyses, except that the significance level to choose significant clusters was one-sided, due to the clear hypothesis.

## Acknowledgements

We thank Christian Keitel and Edwin Robertson for valuable comments on earlier versions of this manuscript. This research was supported by the UK Biotechnology and Biological Sciences Research Council (BBSRC, BB/L027534/1). CK is supported by the European Research Council (ERC-2014-CoG; grant No 646657); JG by the Wellcome Trust (Joint Senior Investigator Grant, No 098433).

## Supplementary figures

**Figure S1.**
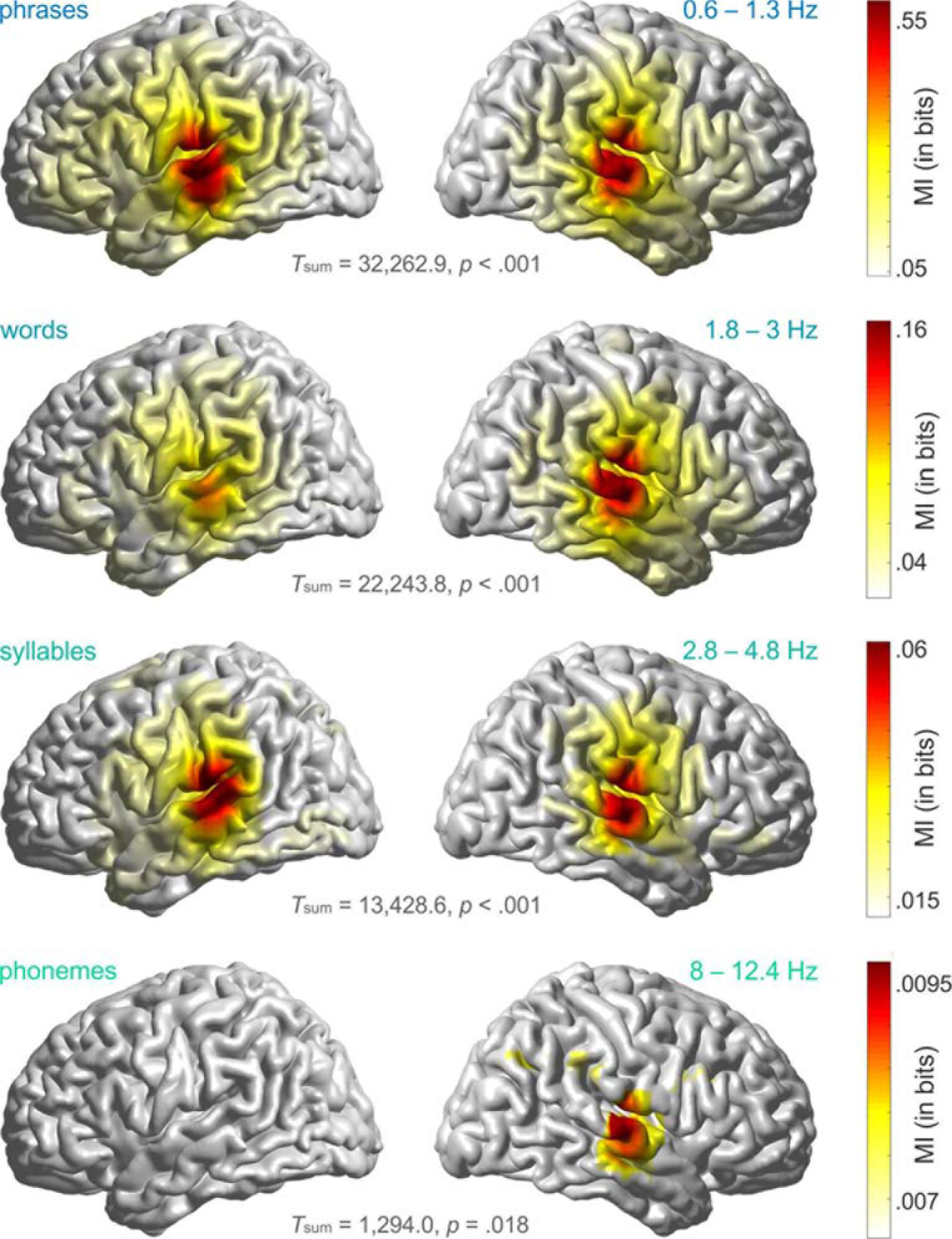
Entrainment in the four stimulus-tailored frequency bands. Shown is significant mutual information for comparison between true MI values and surrogate data (*t*-test, cluster-corrected, all df = 19). The used frequency bands map onto time-scales for phrases (0.6 – 1.3 Hz), words (1.8 – 3 Hz), syllables (2.8 – 4.8 Hz), and phonemes (8 – 12.4 Hz).

**Figure S2.**
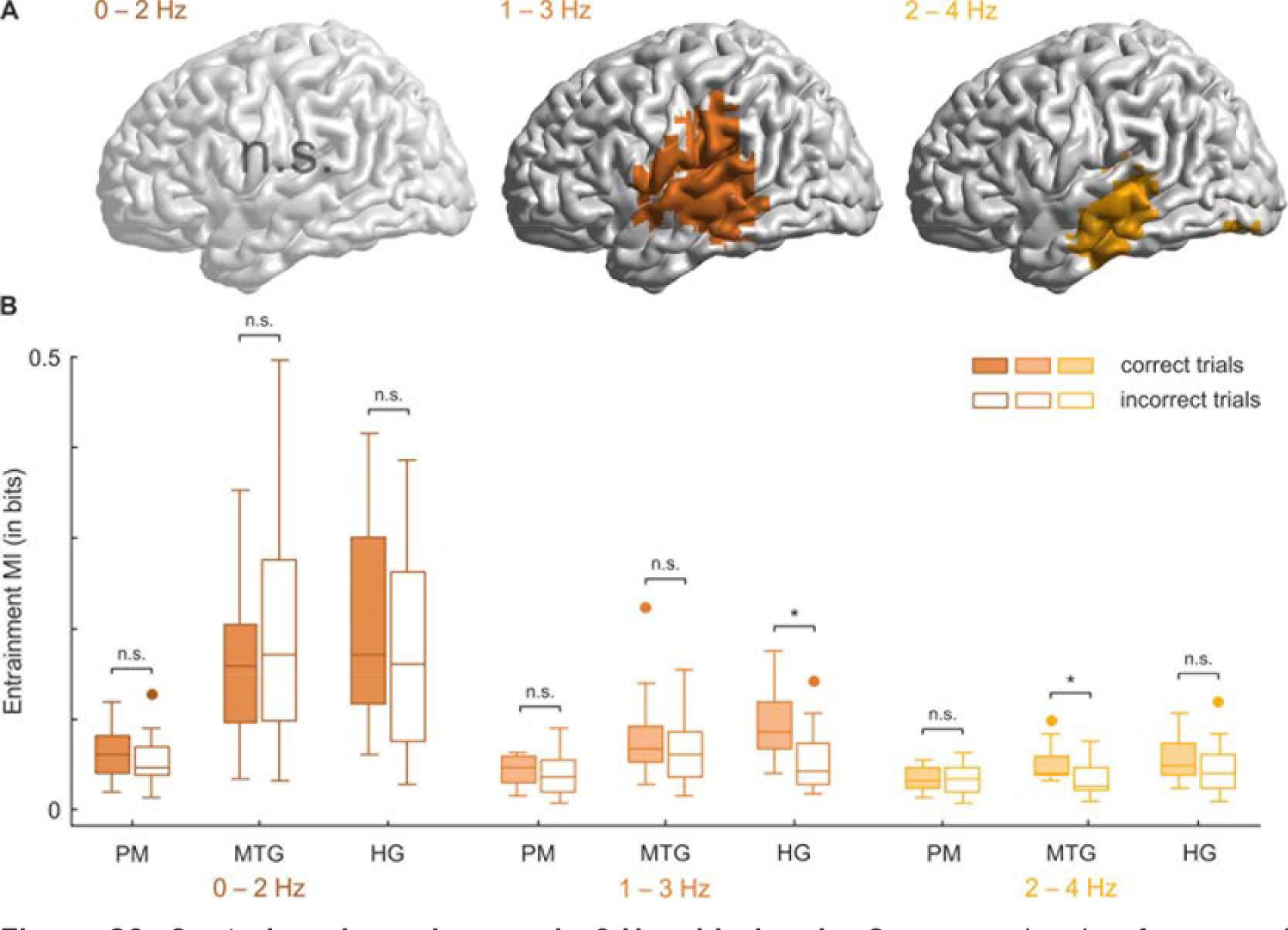
Control analyses in generic 2-Hz wide bands. Seven overlapping frequency bands were analysed (from 0 – 8 Hz, in 2-Hz wide bands, in 1-Hz steps). The first three of these bands are displayed here. A) A significant comprehension effect (larger MI for correctly comprehended than incorrectly comprehended trials) was found at the 1 – 3 Hz scale (*T*_sum_(19) = 1,078.85, *p*_cluster_ = .030) and at the 2 – 4 Hz scale (*T*_sum_(19) = 751.93, *p*_cluster_ = .046). Effects in all other bands were *p* > .11. B) Analysis of the peak grid points that showed the strongest comprehension effect in stimulus-specific bands. Larger MI for correctly than incorrectly comprehended trials is found in Heschl gyrus (HG) in the generic 1 – 3 Hz band (*t*(19) = 4.54, *p*_FDR_ = .002), and in middle temporal gyrus (MTG) in the 2 – 4 Hz band (*t*(19) = 3.38, *p*_FDR_ = .014). All other comparisons are *p* > .08). Notably, the peak grid point in the premotor cortex (PM) does not show a comprehension modulation in any of the generic bands.

**Figure S3.**
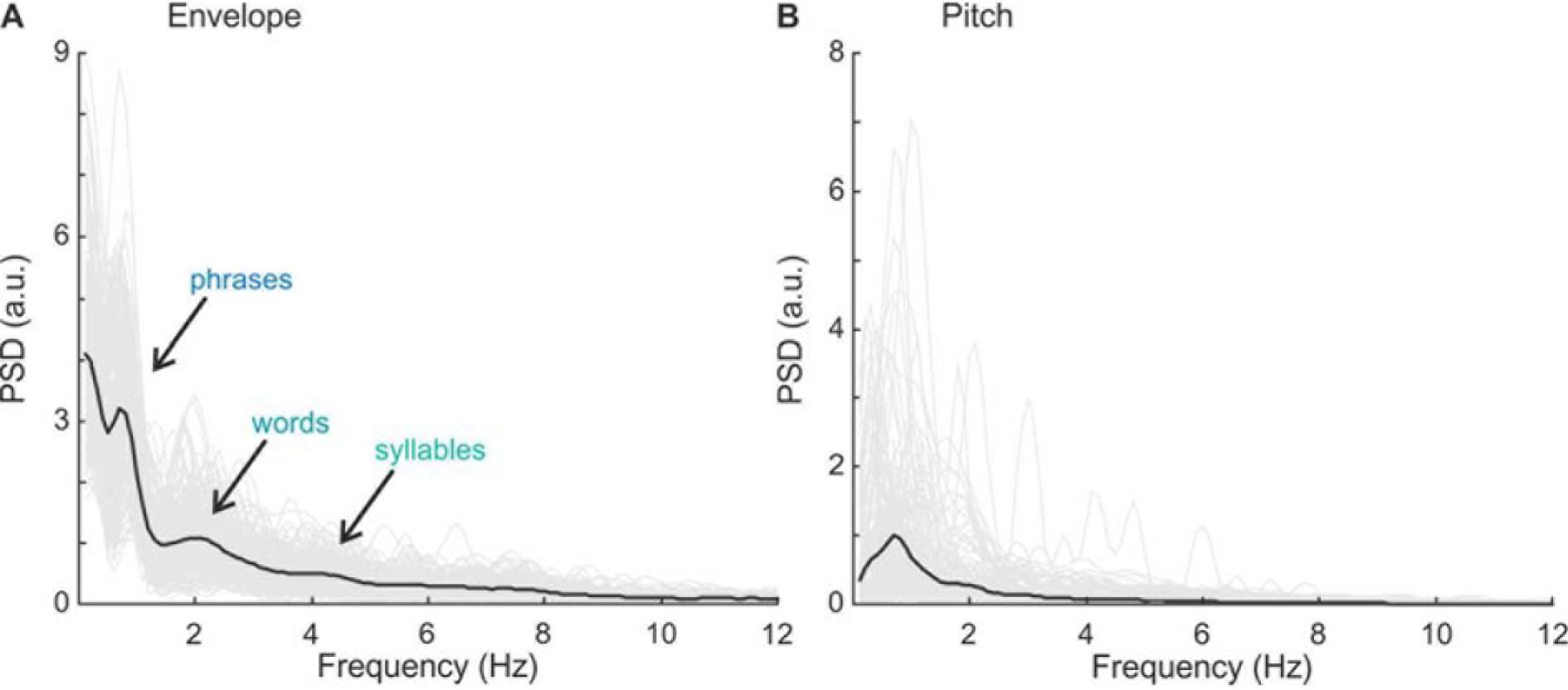
Power spectral density estimates of speech envelope and pitch. Welch’s periodograms are shown for speech envelopes (A) and fundamental frequency (F0-contours/pitch) (B) of all 180 stimulus sentences (thin gray lines) and their average (thick black line), for frequencies between 0.1 and 12 Hz (in 0.1-Hz-steps). For envelope spectra, visible peaks that correspond to rates used in the analysis are marked with arrows (i.e., for phrases, words and, less pronounced, syllables). Fundamental frequency was extracted using Praat [64].

